# *Zea Lip*: An atlas of glycerolipid profiles across leaf development in maize

**DOI:** 10.64898/2026.05.07.723536

**Authors:** Karla A. Juárez Núñez, Guillaume Lobet, Nirwan Tandukar, Eric Jadidzadeh, Asher Pasha, Nicholas J. Provart, James B. Holland, Rubén Rellán-Álvarez, Allison C. Barnes

## Abstract

Lipids are the predominant building blocks of plant membranes and are essential for plant growth and development. They are crucial for survival during times of stress as lipids are involved in multiple signaling pathways, and their relative abundances can change in response to environmental factors. To better characterize the lipid composition of the vital food crop maize, we generated a comprehensive glycerolipid atlas using ultra-high-performance liquid chromatography coupled with quadrupole time-of-flight mass spectrometry. We surveyed the lipid profiles of three different maize genotypes: B73, a temperate inbred; CML312, a subtropical inbred; and Palomero Toluqueño, an open-pollinated variety from the Mexican highlands. We collected leaf samples from 4 developmental stages and 6 leaves. From one growth stage, we also sampled along with three leaf zones: base, center, and tip. The genotype and leaf number were the major drivers of lipid differences. Phosphatidylcholine, lysophosphatidylcholine, and triacylglycerol genotypic differences were particularly high. We generated an eFP browser to be integrated into the maize genome browser, as well as a separate web interface to easily browse and compare lipid levels across tissues and genotypes, available at https://bar.utoronto.ca/~dev/efp_maize_lipid/cgi-bin/efpWeb.cgi and https://rrellan.shinyapps.io/Zea-Lip/respectively.

**SIGNIFICANCE STATEMENT:** This work creates a spatial map of lipids in maize leaves across four growth stages for three genotypes: a lowland, a sub-tropical, and a highland. The resources generated here will directly benefit both the maize and lipid communities by creating a large dataset that can be used to generate new hypotheses in understanding lipid metabolism and environmental responses in maize.

## INTRODUCTION

Lipids are essential components of plant tissues, involved in everything from growth and development to stress tolerance and hormone signaling mechanisms (Quinn and Williams, 1979; Wang, 2004; Moellering and Benning, 2011; Li and Yu, 2018). Glycerolipids make up the vast majority of lipids in plant cells. They contain a glycerol backbone with an ester bond to fatty acid tails; they can be separated into two subgroups: galactolipids and phospholipids. Galactolipids, such as monogalactosyldiacylglycerol (MGDG) and digalactosyldiacylglycerol (DGDG), are specific to plants and algae and are the most abundant lipids in nature (Lee, 2000). Their ratios are crucial for stabilizing photosynthesis protein components and giving chloroplasts their shape (Mazur *et al*., 2019; Seiwert *et al*., 2017; Yu *et al*., 2020). Phospholipids can have different polar head groups and readily shift between them. Phospholipases A, B, C, and D shuttle existing phospholipids into variations of DAG and phospho-headgroups, particularly during times of stress. Phospholipids are the most prevalent lipids outside the chloroplast.

Phospholipid metabolism is complex but can be interconnected through phosphatidic acid (PA) and free or activated diacylglycerol (DAG). The most abundant phospholipids in plants are phosphatidylcholine (PC) and phosphatidylethanolamine (PE), which share overlapping biosynthetic pathways and are interconvertible. Phosphatidylserine (PS) is also linked to PE synthesis. PC, PE, and PS all have a free DAG substrate, which also connects them to galactolipid synthesis. Other phospholipids, such as phosphatidylglycerol (PG) and phosphatidylinositol (PI), share the activated CDP-DAG substrate. PI is an important signaling lipid, while PG and cardiolipin (CL), which consists of two PG molecules, are important for chloroplast and mitochondrial function, respectively. The composition of the membrane, particularly the shape of the lipids that form the bilayer, will affect its properties and functionality. The lipid bilayer was identified 100 years ago (Gorter and Grendel, 1925). However, plants’ specific bilayer variations, which serve as adaptations to different environments, are still being uncovered.

One challenge in studying lipid composition in plants is the complexity of the mixture. Plant lipid profiles depend highly on the tissue sampled, the environment, and even the time of day the sample was taken (Kenchanmane Raju *et al*., 2024; Nakamura *et al*., 2014; Kenchanmane Raju *et al*., 2018). High-throughput lipidomics of plants using mass spectrometry began early in the 21st century (Welti *et al*., 2002; Wang, 2004). With the development of new methods, the number of samples that could be processed increased. In conjunction, a decrease in the amount of time required per sample and an increase in the power to detect, separate, and identify different lipid species occurred. Lipidomes of many types of plants, including model, agricultural, and ornamental plants, are now available (Kehelpannala *et al*., 2021; Matsuzawa *et al*., 2021; Deng *et al*., 2025; Narayanan *et al*., 2016; Zhu *et al*., 2022). However, because of environmental, developmental stage, tissue, and genotype variation, each lipidome is essentially a snapshot of lipids at that single point in time.

Maize leaf lipids have historically been studied with respect to their effects on chloroplasts, anatomy, and their relationship to photosynthetic ability (Leech *et al*., 1973; Hawke *et al*., 1974; Bolton and Harwood, 1978; Leese and Leech, 1976). Much of this work looked at the differences between mesophyll and bundle sheath chloroplasts and the lipid differentiation from proplastid to chloroplast. Leech *et al*. state, “The glycolipids are quantitatively the most important lipids at all stages of cellular differentiation in the maize leaf sections.” (1973). These studies observed increases in PG, MGDG, and DGDG as cells developed from the base of the leaf to the tip, most likely due to an increase in membrane content associated with the greater number of plastids in older tissue.

Here, we sought to compare the leaf lipidomes of cells from different leaves at different developmental stages from three maize genotypes of diverse origins. Because these genotypes are inherently tolerant to different stresses and adapted to distinct environmental conditions, we hypothesized that their lipid composition profiles would differ even under controlled growth conditions. B73 is a temperate, inbred line and a good historical representative of the stiff-stalk heterotic group, CML312 is a subtropical inbred line adapted to a wide variety of environments in Mexico, and Palomero Toluqueño (PT) is an open-pollinated variety from the central Mexican highlands (2597 masl).

In *Zea mays* and other monocots, leaf differentiation occurs basipetally, with the oldest cells at the tip and the youngest at the base (Sharman, 1942; Nelson and Dengler, 1997). It has also been reported that there is a contrasting gene expression pattern between the tips and bases of maize leaves at the same or different developmental stages (Sekhon *et al*., 2011). Thus, we sampled tissue from three leaf zones: base, center, and tip. We also sampled 4 growth stages and 6 leaves, numbered according to their order of appearance in development, using greenhouse-grown plants. We selected these leaf numbers and zones to sample maize leaf development along its two principal axes. Successive leaf positions (leaves 4, 5, 7, 9, 11, and 13) mature sequentially, so harvesting several leaves at one date captures a range of developmental stages at once, while the base–center–tip axis captures the basipetal differentiation gradient within a single leaf. Because much previous maize lipidomic work has focused on seedlings and young plants, we deliberately extended sampling across a wider range of plant maturity and leaf positions. Indexing development by leaf number and within-leaf zone, rather than by chronological age alone, allows samples to be compared at equivalent developmental positions across genotypes that differ in absolute growth rate Using Ultra-High-Performance Liquid Chromatography coupled to a Quadrupole Time of Flight Mass Spectrometry (UHPLC-QTOF MS/MS), we identified 149 molecular lipid species within 14 lipid classes. Because our aim was to build a broad reference atlas spanning genotype, developmental stage, leaf position, and leaf zone, we prioritized sampling breadth deep replication within any single combination of factors. The resulting dataset is intended as a hypothesis-generating community resource; measurements corresponding to individual cells of the design that are represented by a single plant should be interpreted accordingly, whereas the statistical inferences reported here draw their power from the full structure of the design across all samples. This study provides unique insight into lipid distribution during maize leaf development and a database of lipid composition profiles, available in a convenient format for the plant science community to access.

## MATERIALS AND METHODS

### Plant growth and tissue collection

B73, a temperate inbred; CML312, a subtropical inbred; and Palomero Toluqueño (PT), an open-pollinated traditional variety from the Mexican highlands, were grown in a greenhouse at an average temperature of 23 °C to 30 °C (night/day) under an average of 13 and a half hours of light (6:00 am to 19:20 pm). All plants were watered with untreated water and were fertilized with 14 g of Osmocote every 18 days. Plant samples were collected between 10:00 am and 12:00 pm.

Germination was initiated by soaking the seeds in distilled water in Falcon tubes for 24 hours. After germination, plant growth was recorded daily, including the number of emerging and fully expanded leaves per plant. The coleoptile was considered the first leaf. Plant growth and shoot/root ratios of these genotypes under these conditions are shown in Figures S2 and S3.

Seeds from all genotypes were sown in four blocks, with an approximately 10-day difference between the sowing dates. From all plants, 50 mg of fresh blade tissue was collected 58 days after the first sowing date, thereby sampling multiple developmental stages (28, 40, 50, and 58-day-old plants) at once (Figure 1A, Table S1). In total, 50 plants were sampled yielding a total of 151 samples. All samples were taken along the length of the leaf blade and excluded the midrib, ligule, and auricles. From the 58-day plants, leaf 9 was also sampled in three different zones: tip, center, and base (Figure 1B). Tissue was collected from the rest of the planting dates along the leaf blade. The midrib, the ligule, and the auricles were always excluded while sampling. The tissue collected was immediately frozen in liquid nitrogen and stored at -80 °C until lipid extraction.

**Figure 1.**
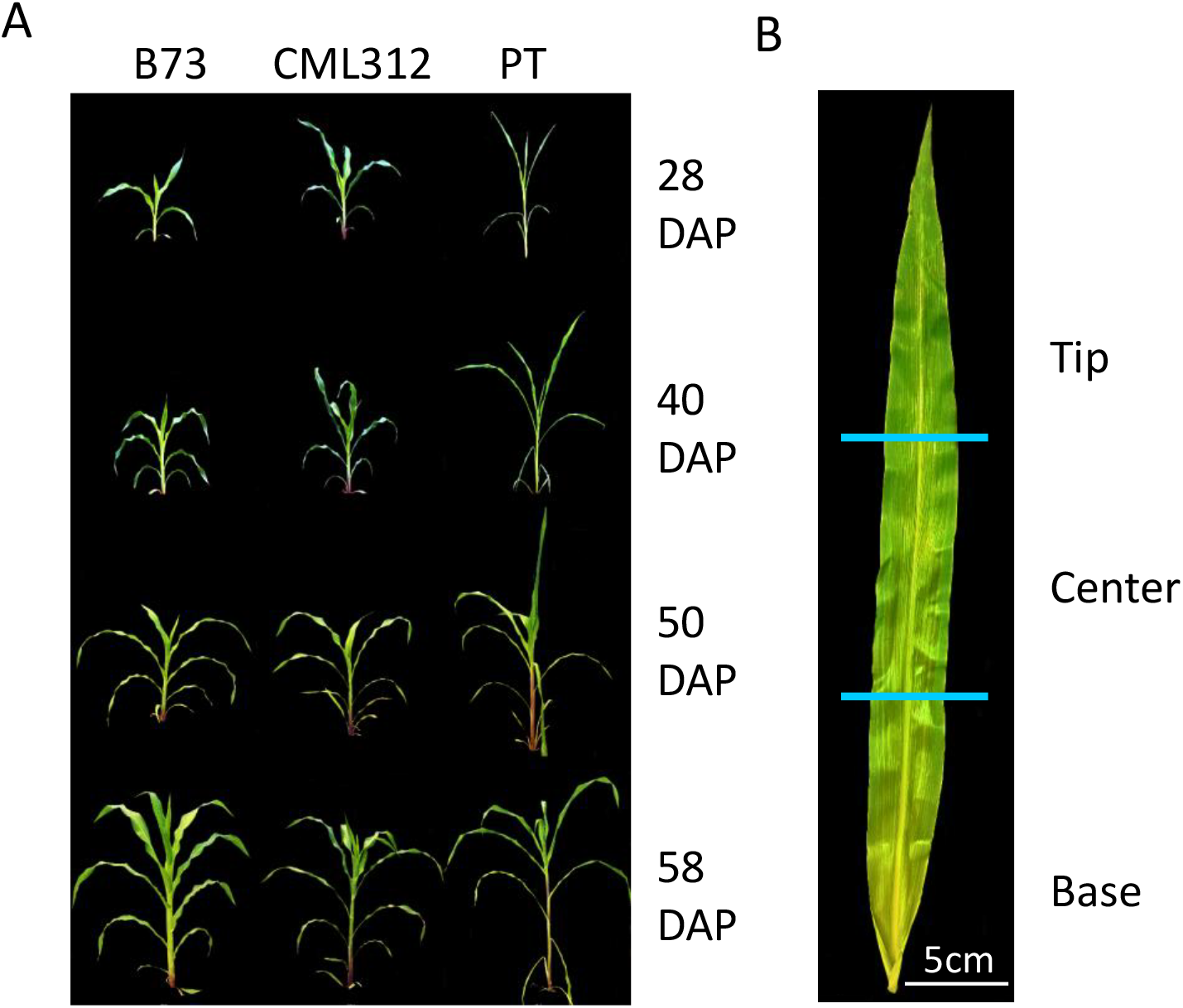
Maize genotypes and growth stages at collection. **A)** B73, CML312, and PT genotypes shown at each sampling date. They were sown in four different blocks, having a 10-day difference between the four sowing dates so that all growth stages could be sampled in one day to reduce the variation in samples. **B)** Leaf zones included in lipid profiling from the first planting date (58 DAP). The tissue was collected along a 5 cm distance for the base, starting 1 cm above the auricles and ligules to exclude them. The tip was sampled from 2 cm below the tip, along a 5 cm distance. Finally, the center in short leaves was sampled using a separation of 2 cm distance from the end of the base and the tip. For longer leaves, distances of 10 and 20 cm between the base and tip to center were used.

### Lipid extraction and UHPLC-QTOF MS/MS lipid profiling

Fifty mg of leaf tissue was collected using a leaf puncher and snap-frozen in liquid nitrogen. Frozen material was homogenized in a tissue grinder Retsch (Haan, Germany) for 40 seconds at a frequency of 30 Hz. After grinding, samples were immediately extracted.

We performed lipid extraction as reported by Matyash and collaborators (Matyash *et al*., 2008). First, 225 μL of cold methanol (MeOH) was added to each sample. Blank samples were prepared in MeOH using a Quality Control (QC) mix ( Table S2). Each sample was vortexed for 10 seconds, while the remaining materials remained on ice. Then 750 μL of cold methyl tert-butyl ether (MTBE) was added. MTBE added to blank samples also contained 22:1 cholesterol ester as an internal standard Table S2). Each sample was vortexed for 10 seconds, then shaken for 6 minutes at 4°C in an orbital mixer. 188 μL of LC/MS-grade water at room temperature was added, and the samples were vortexed for 20 seconds.

Samples were then centrifuged for 2 min at 14000 rcf (12300 rpm), and 700 μL of the upper organic phase supernatant was collected. The supernatant was split into two aliquots of 350 μL, one for lipid profiling and the other for the preparation of pools to be used with the lipid profiling. Finally, samples were dried using a speed-vacuum concentrator.

Dry samples were resuspended in 110 μL of MeOH-Toluene 90:10 containing the internal LC/MS standard CUDA (12-(3-cyclohexylureido)dodecanoic acid) at 50 ng/mL Samples were vortexed at low speed for 20 s and then sonicated at room temperature for 5 min. Aliquots of 50 µL per sample were transferred for mass spectrometry.

Samples were analyzed on UHPLC-QTOF MS/MS Agilent 1290 and Agilent 6530, respectively. The UHPLC used a Waters Acquity charged surface hybrid (CSH) C18 2.1x100 mm, 1.7 μm column coupled to a VanGuard pre-column. The column was initially purged for 5 min. Six “no sample injections” were injected at the beginning of each run to condition the column, followed by ten samples, one pool (made from a mix of the second aliquot of all the samples combined), and one blank.

To identify the appropriate electrospray ionization mode, we analyzed MS1 spectra in both positive and negative modes for 8 B73, 8 PT, and 10 recombinant inbred line (RIL) samples derived from the cross between B73 and PT (Barnes *et al*., 2022). This optimization step identified 24 unique lipid species across six classes (CE, DG, LPC, PC, PE, TG) under positive mode, versus 16 species (FA, PC) under negative mode, with no overlap in species identified between modes (Table S4). Table S4 reports this preliminary mode-comparison dataset only and is independent of the developmental-stage sampling design used in the main study; the leaf-number/leaf-zone series (leaves 4, 5, 7, 9, 11, 13; base, center, tip zones) described below does not apply to these samples. Sample labels in Table S4 follow the format [Replicate] [Genotype/Line] [Block] [Tissue], where the leading numeral (1, 2) In the “Label” row, the leading numeral (1, 2) denotes two biological replicates per genotype, the letters A - D denote the block-number/ position in the greenhouse where each plant was growing, the genotype or RIL identifier follows and “L” indicates leaf tissue; a full legend is provided in Table S4. Based on this optimization, the positive mode was selected for analysis.

Injections of 1.67 μL per sample were used. The running time per sample was 15 min. Mobile phase “A” consisted of 60:40 acetonitrile:water, 10 mM ammonium formate, and 0.1% formic acid. Mobile phase “B” consisted of 90:10 isopropanol:acetonitrile, 10 mM ammonium formate, and 0.1% formic acid. The flow rate was maintained at 0.6 mL/min, and the column compartment was maintained at 65 °C. Initial conditions were 15% B; the gradient increased uniformly to 100%. At 12.10 min, the mobile phase composition returned to initial conditions. For the source parameters, ESI gas temperature was set at 325 °C, nebulizer pressure at 35 psig, gas flow at 11 L/min, capillary voltage at 3500 V, nozzle voltage at 1000 V, and MS TOF fragmenter and skimmer at 120 and 65 V, respectively. Under the acquisition parameters, a mass range of 60-1700 m/z was set, with a detection window of 100 ppm and a minimum height count of 1000.

### Data processing and analysis

Corrected retention time (RT) was calculated using Agilent MassHunter Qualitative Analysis® B.06.00 version and Microsoft Excel. Agilent MassHunter Qualitative Analysis was used to extract ion chromatograms (EICs) of the internal standards within the run. The highest intensity point for each EIC was then used to determine the current RT. The reported RT for internal standards was compared to the current RT, and a polynomial regression was performed. The subsequent equation was used to calculate adjusted retention times for a 501 lipid MS1 m/z-rt library ( Table S3).

To check for false positives, 36 samples from the study were run across four m/z ranges: 120-300, 300-600, 600-900, and 900-1200. MSDIAL (Tsugawa *et al*., 2015) was used to determine how many fragments of each identified lipid matched spectra from the Lipidblast library (Kind *et al*., 2013). This analysis enabled the addition of 339 new annotations to the mz-rt library, bringing the total to 840 lipid species. This library was used to reprocess the spectra using MSDIAL and MS-FLO (DeFelice *et al*., 2017). The identification of lipids was based on two approaches: an internal library with *in silico* MS/MS comparisons and a user-supplied library with corrected RTs (Table S5).

Raw data files were converted from .d to .abf format using the Reifycs Abf converter (https://www.reifycs.com/AbfConverter/) and imported into MSDIAL version 3.40 (Tsugawa *et al*., 2015). The MSDIAL alignment results were filtered to remove any compounds with an intensity less than 10 times that of the blank. Filtered data were normalized to quality control samples by Systematic Error Removal using Random Forest (SERRF) (Fan *et al*., 2019). Normalized features with a coefficient of variation (CV) of 30% or greater across the pools were removed. To curate data on duplicate features, isotopes, and ion adducts, we used MS-FLO (DeFelice *et al*., 2017). Individual lipid intensities were further normalized by dividing each by the total ion count (mTIC) of all known and quantified lipids in each chromatographic run.

The separation of maize lipids occurred from 0.5 to 12 min (Figure S1), with some lipid classes overlapping in elution time. From 0.5 to 1.1 min, acylcarnitine (AC) species eluted from the column, followed by lysophosphatidylcholine (LPC) species from minute 1 to 2.3 min. Sphingomyelins (SM) eluted between 3.3 and 6.7 min, and phosphatidylglycerols (PG) species eluted in a period from 3.2 to 5 min. Phosphoethanolamines (PE) eluted from 3.3 to 7.8 min, and digalactosyldiacylglycerols (DGDG) between 3.8 and 8.1 min. Species of monogalactosyldiacylglycerol (MGDG) eluted between 3.92 and 6.5 min, and from 4.3 to 7.21 min, phosphatidylcholine (PC) species were identified. Sulfoquinovosyldiacylglycerols (SQDG) eluted from 4 to 4.81 min. From 4.5 to 7.1 min, glucosylceramides (GlcCER) eluted while diacylglycerols (DG) were found between 5 and 6.4 min. From 5.2 to 5.8 min, ceramides (CER) eluted. The most abundant lipid class identified was the triacylglycerols (TG), with 42 TG species that eluted between 6.5 and 11.4 min. The only cholesterol ester (CE) species identified eluted at 10.33 min.

### Statistical Analysis

#### Effect of the leaf zone

To determine whether the leaf zone caused significant variation in lipid profiles while controlling for other experimental factors, we used the stepAIC function (Venables and Ripley, 2002) to sequentially fit factors into a linear model for each compound. Using data from only the first planting date, we tested the effects of genotype, zone, leaf, and their interactions. For each compound, this analysis identifies a linear model in which all factors contribute to improved model fit, based on Akaike’s Information Criterion (AIC) (Cavanaugh and Neath, 2019). The maximum model that could be fit was *Y* ∼ 1 + genotype + zone + leaf + genotype:leaf + genotype:zone + leaf:zone + genotype:leaf:zone + residual, where *Y* was the normalized value for one lipid in a particular sample, 1 refers to the intercept, and colon symbols indicate interactions between factors. Model selection was conducted in the forward direction.

To compare the importance of different model factors across all compounds, we computed the frequency with which each model factor was selected by stepAIC. To examine the variance contributed by each factor, we compared the distribution of the proportion of sums of squares across compounds.

Compounds that had significant variance caused by zone or zone and genotype interactions were then examined further by fitting the following linear model: *Y* ∼ 1+ zone + genotype:zone + leaf:zone + genotype:leaf:zone, and then estimating marginal means for genotype, zone, and leaf combinations using the emmeans package (Lenth, 2025). The results were then summed to create total values for each lipid class.

#### Effects of genotype, developmental stage, and leaf number

To determine the relative importance of genotype, developmental stage, and leaf number on lipid profiles, we used the data from all planting dates. Again, we used stepAIC (Venables and Ripley, 2002) to test each of the following model factors for inclusion in a final model: genotype, leaf number, developmental stage, and all their two- and three-way interactions. We also included the planting date factor, but without interactions. We then calculated the proportion of the sums of squares for each factor retained for each lipid. We estimated mean quantities of each lipid for each combination of leaf and genotype by fitting the following linear model: *Y* ∼ 1 + genotype + leaf + genotype:leaf + planting date + developmental stage. Adjusted means for genotype and leaf combinations were estimated using the emmeans package (Lenth, 2025). We estimated the frequency with which a model factor was chosen across lipids.

#### Principal Component Analysis

Principal component analysis was applied to the mTIC-normalized lipid values using the prcomp function in the stats package in R (R Core Team, 2024).

#### Visualizations

All visualizations were made with the ggplot2 package (Wickham, 2016) in R.

## RESULTS AND DISCUSSION

### Separation, annotation, and identification of maize leaf lipid classes

To achieve a thorough profile of the maize leaf lipidome, we selected three maize genotypes with origins in diverse environments: B73, Palomero Toluqueño (PT), and CML312. B73 is a temperate inbred, CML312 is a subtropical inbred, and PT is an open-pollinated variety from the highlands of Mexico. At a single growth stage, we harvested tissue from three leaf zones: base, center, and tip. Tissue from all along the blade was also sampled at four separate growth stages. As the three genotypes are adapted to very different environmental conditions, especially contrasting temperatures, we expected the lipid profiles to differ, even in a greenhouse environment. Because leaf development occurs basipetally, we also expected to identify variations across leaf zones.

Extracted lipids were analyzed using UHPLC-MS/MS. Over 19,000 mass features were identified with an MS1-MS2 m/z-rt lipid library and LipidBlast (Kind *et al*., 2013), resulting in 149 lipids from 14 lipid classes (Table 1). We annotated the lipids following the LIPID MAPS classification system (Fahy *et al*., 2009; Liebisch *et al*., 2020), as a standardized shorthand is important for cross-species analysis and discussion.

**Table 1.** Lipid classes identified and the number of lipid species per class.

| Lipid Class | Subcategory and Number of Species Identified |
| --- | --- |
| Fatty Acyls (FA) | *Fatty Acylcarnitine [FA0707]*<br>□ Acylcarnitine (AC) - 1 |
| Sterol Lipids (ST) | *Sterol Ester [T0102] -*<br>□ Cholesterol esters (CE) - 2 |
| Sphingolipids (SP) | *Ceramides [SP0201]*<br>□ Ceramides (CER) - 2<br>*Simple Glycosphingolipids [SP0501]* |
|  | □ Glucosylceramides (GlcCER) - 6 |
| Glycerolipids (GL) | *Triacylglycerols [GL0301]*<br>□ Triacylglycerols (TG) - 42<br>*Diacylglycerols [GL0201]*<br>□ Diacylglycerols (DG) - 9<br>*Glycosyldiacylglycerols [GL0501]*<br>□ Monogalactosyldiacylglycerols (MGDG) - 12<br>□ Digalactosyldiacylglycerols (DGDG) - 19<br>□ Sulfoquinovosyldiacylglycerols (SQDG) - 9 |
| Glycerophospholipids (GP) | *Monoacylglycerophosphocholines [GP0105]*<br>□ Lysophosphatidylcholines (LPC) - 6<br>*Diacylglycerophosphocholines [GP0101]*<br>□ Phosphatidylcholines (PC) - 21<br>*Diacylglycerophosphoethanolamines [GP0201]*<br>□ Phosphoethanolamine (PE) - 11<br>*Diacylglycerophosphoglycerols [GP0401]*<br>□ Phosphatidylglycerols (PG) - 8<br>*Diacylglycerophosphoserines [GP0301]*<br>□ Phosphatidylserine (PS) - 1 |

Lipid classes and subcategories were assigned according to the LIPID MAPS lipid classification system (Fahy *et al*., 2009). The number code assigned by LIPID MAPS to the lipid subcategories is indicated with brackets, while the abbreviation used for each lipid within each subcategory is indicated within parentheses.

### Maize lipid profiles differ by genotype and leaf number

After quantifying as many lipid compounds as possible, principal component analysis was used to identify important trends in the data. Principal Component 1 (PC1) accounted for 31.7%, and Principal Component 2 (PC2) accounted for 16.8% of the variance across all lipid species (Figure 2A, B). Clear, predictable lipid relationships were observed. For instance, PC and lyso-phosphatidylcholine (LPC) were clearly separated on both PC1 and PC2 (Figure 2A). These two lipids differ by one fatty acid tail, and their separation on the PC plot represents their negative correlation driven by the production of LPC from PC. MGDG 36:6 and DGDG 36:6, two lipids that differ simply by one sugar moiety on the headgroup, clustered together, reflecting the positive correlation between their abundances. In general, the TGs clustered together. PE, PC, PG, MGDG, and DGDG also clustered together. These lipids regularly interconvert and shuttle between the endoplasmic reticulum and the chloroplast, as lipid metabolism in plants is intricately connected (for reviews on lipid metabolism and transport, see (LaBrant *et al*., 2018; Lavell and Benning, 2019). MGDG and DGDG differ by just one galactose moiety on the head group and are made in the chloroplast along with PG. PC can be generated from PE by three methylations (Bolognese and McGraw, 2000). All originate from DG lipid backbones that enter multiple pathways. To examine relationships between lipid groups as a whole, we repeated the principal component analysis using the sums of each lipid class (Figure S3). These results were congruent with the individual lipid species analysis.

**Figure 2.**
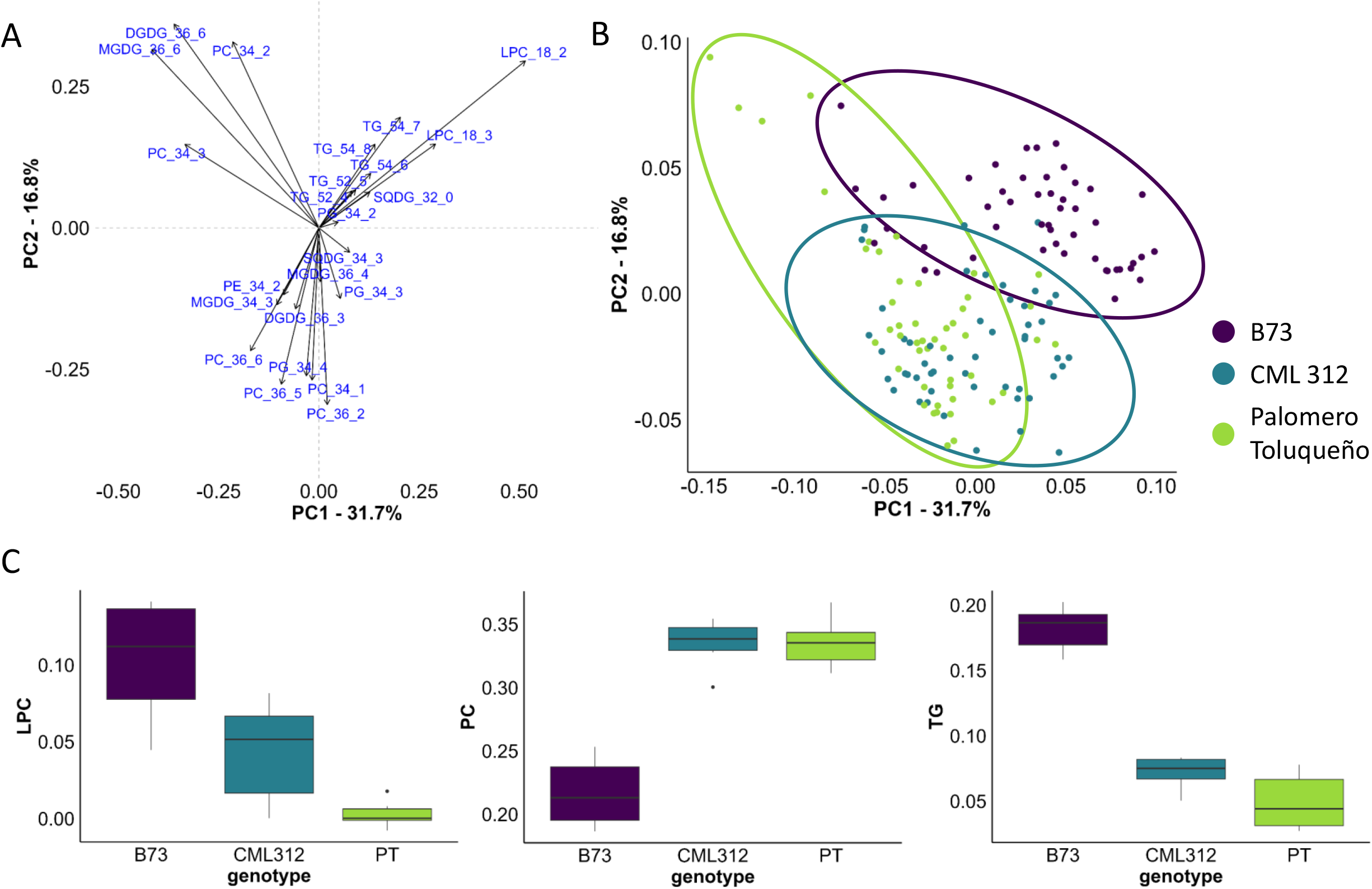
Phosphatidylcholine, lysophosphatidylcholine, and triacyglycerol drive genotype separation in principal component analysis. **A)** Principal component analysis using scaled values of all lipids measured. Principal component 1 explains 31.7% of the variation and principal component 2 explains 16.8% of variation. Any lipid with an absolute value of variance of less than 0.04 is excluded from the plot for viewing simplicity. **B)** Principle components 1 and 2 plotted as a percentage of total variation and colored by genotype. **C)** Sums of lysophosphatidylcholine, phosphatidylcholine, and triacylglycerol for each of the three genotypes, demonstrating the differences between them. B73 and PT are generally opposites, with CML312 as an intermediate phenotype.

To further explain the variance, we examined the genotype, leaf number, and developmental stage of the samples and found that genotypes are the most important factor separating the samples along PC1 and PC2 (Figure 2B). B73 and PT exhibit opposing patterns, with CML312 clustering between them. This variation corresponds to the origins of these genotypes: PT is adapted to highlands, B73 to US midwestern temperate environments, and CML 312 to subtropical environments. This pattern was also observed when we examined lipid sums and some relevant ratios (Figure 2C). For example, PC is high in PT plants and low in B73, with an intermediate phenotype in CML312. LPC is low in PT plants, high in B73, and intermediate in CML312 (sums of all lipid classes available in Figure S4).

The particularly strong genotypic variation in PC and LPC is consistent with the activity of the HIGH PHOSPHATIDYLCHOLINE 1 (HPC1) locus, which we previously characterized in maize and its wild relatives (Barnes et al., 2022). HPC1 encodes a phospholipase A1–type enzyme that hydrolyzes PC to LPC; a teosinte mexicana–derived introgression prevalent in highland germplasm reduces HPC1 activity, resulting in higher PC and lower LPC. The lipid profiles reported here mirror this relationship: the highland-adapted Palomero Toluqueño accumulates high PC and low LPC, the temperate B73 shows the opposite, and the subtropical CML312 is intermediate (Figure 2C). Because PC and LPC are directly interconverted by this phospholipase reaction, genotypic differences in PC/LPC balance likely reflect variation in phospholipase activity associated with highland adaptation. The comparably strong variation in triacylglycerols is consistent with the role of TAG as a transient reservoir for fatty acids released during membrane remodeling and turnover, a process expected to differ among genotypes adapted to contrasting thermal environments and one that we find is further structured by leaf number and developmental stage (Figure 3).

**Figure 3.**
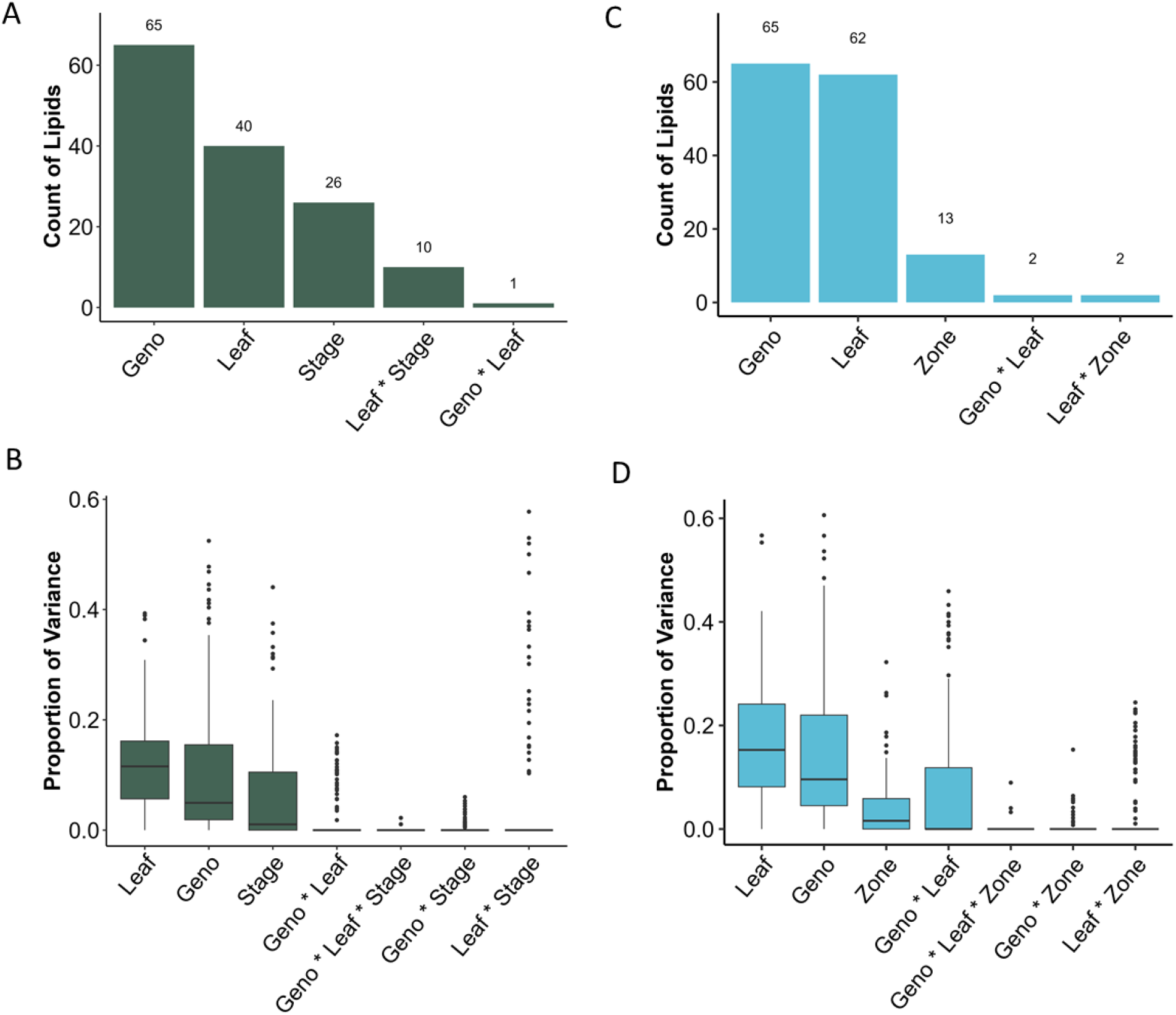
Genotype and leaf number are the two most important factors in determining lipid differences. Using data across all planting dates, a stepwise regression model was used to identify the experimental factors that contributed significantly to the variance. The most significant factor for each lipid is summed **(A)**, and the distribution across lipids of the proportion of the variance accounted for by each factor was calculated **(B)**. Using data from just the 58 DAP samples, we removed the developmental stage and added the leaf zone to identify the significant factors for each lipid. The most significant factor for each lipid is summed **(C)**, and the distribution across lipids of the proportion of the variance accounted for by each factor **(D)**.

To measure the relative importance of the different experimental factors (leaf number, genotype, developmental stage, and their two- and three-way interactions) on variation in each lipid species, we performed forward regression modeling for each lipid, using AIC as the criterion for including factors in the final model. This model sequentially tests each factor or interaction, adding the one associated with the most variation at each step, and stops when no additional model terms explain further variation. Genotype or leaf number was the most important factor or factor interaction (explaining the most variation among samples) for most lipids (Figure 3A). Because genotype and leaf were the most significant factors for so many lipids, there was no discernible pattern to the lipids that attributed their primary differences to these two factors. However, some lipid relationships were observed that were most strongly affected by other experimental factors. For example, TG species were the only lipids whose most significant factor was the interaction of leaf number and genotype or leaf number and developmental stage. The developmental stage main effect was the most important factor for a nearly even mix of TG and galacto/sulfolipids, with a few phospholipids. Galactolipids have a crucial function in photosynthesis, so it is not surprising that their variation can be strongly affected by the developmental stage of the plant.

For each experimental factor and its interactions, we also measured the number of lipids that significantly affected (Figure S5A). Genotype and leaf number were the two most commonly significant factors across all lipids, included in the final model for 79% and 77% of the 149 lipids, respectively. The developmental stage was significant for 50% of the lipids, and the two-way and three-way interactions were significant for 23% or less of the lipids (Figure S5A). A final measure of experimental factor importance was the proportion of sums of squares they account for, and by this criterion, it is clear that genotype, leaf, and developmental stage drive most of the variance for most lipids (Figure 3B). While the proportion of variance explained by the leaf-by-stage interaction is usually zero, it occasionally accounts for a large portion of the variation. We further explored which lipids are in this category. This category was overwhelmingly composed of TG species. Vegetative tissue typically does not accumulate large amounts of TG, less than 0.1% dry weight (Xu and Shanklin, 2016). TG is known to be rapidly turned over during leaf senescence and may serve as a transient buffer for released fatty acids, which could explain the correlation between the number of leaves sampled and the plant’s age (Troncoso-Ponce *et al*., 2013).

### Lipid Changes Across Leaf Zones

From the same genotypes, we collected leaf tissue from a single planting date, 58 days before sampling, from multiple leaves, and 3 different leaf zones: the base, the center, and the tip. Leaf number, genotype, zone sampled, and the interactions between these variables were modeled to examine if specific portions of the leaf tissue yielded different lipid content (Figure 3C). Genotype or leaf number was the most significant factor for many lipids, with zone being most important for only 13 of 149 lipids. Genotype and leaf were also significant factors influencing variation in most lipids, but leaf zone was significant for 84 lipids, and the interaction terms were also common significant contributors to lipid variation (Figure S5B). Across lipids, the four factors of leaf, genotype, zone, and genotype by leaf interaction had the most important contributions to total variance (Figure 3D).

We then examined which lipids are influenced by the leaf zone. MGDG, DGDG, PG, TG, and PC all had at least one lipid species for which zone was the most significant factor. Unlike previous reports, we did not observe the highest amount of photosynthetic lipids such as MGDG, DGDG, and PG (Figure 4) in the tip of the leaves consistently across genotypes (Leech *et al*., 1973; Hawke *et al*., 1974; Bolton and Harwood, 1978; Leese and Leech, 1976). PT maintained consistent levels of each of these lipid classes across leaf zones, but CML312 and B73 had a trend of lower levels of MGDG and DGDG in tips compared to leaf bases (Figures 4A and 4B). The levels of PG increased in tips vs. bases for CML312 and B73, consistent with previous reports; again, however, PT showed consistent levels of PG across zones (Figure 4C). The differences from previously reported studies could be due to many factors, including plant age, genotypes used, growth conditions, and the definition of the leaf base, among others. Broadly speaking, the base and the tip seem to be the two zones with the most different lipid profiles. In maize leaves, the oldest tissue is at the tip and the newest at the base; thus, these samples represent two contrasting developmental stages. The difference in tissue age would affect the amounts and types of lipids required for cellular development or differentiation. For example, the base of the leaf receives less sunlight than the center and tip of the leaf. In the shade, chloroplast thylakoid stacks increase in number and thickness to try and capture as much light as possible (Melis and Harvey, 1981; Lichtenthaler, 1984). This could explain the observation of more MGDG at the base of the leaf, as MGDG is the primary lipid in thylakoids (Block *et al*., 1983).

**Figure 4.**
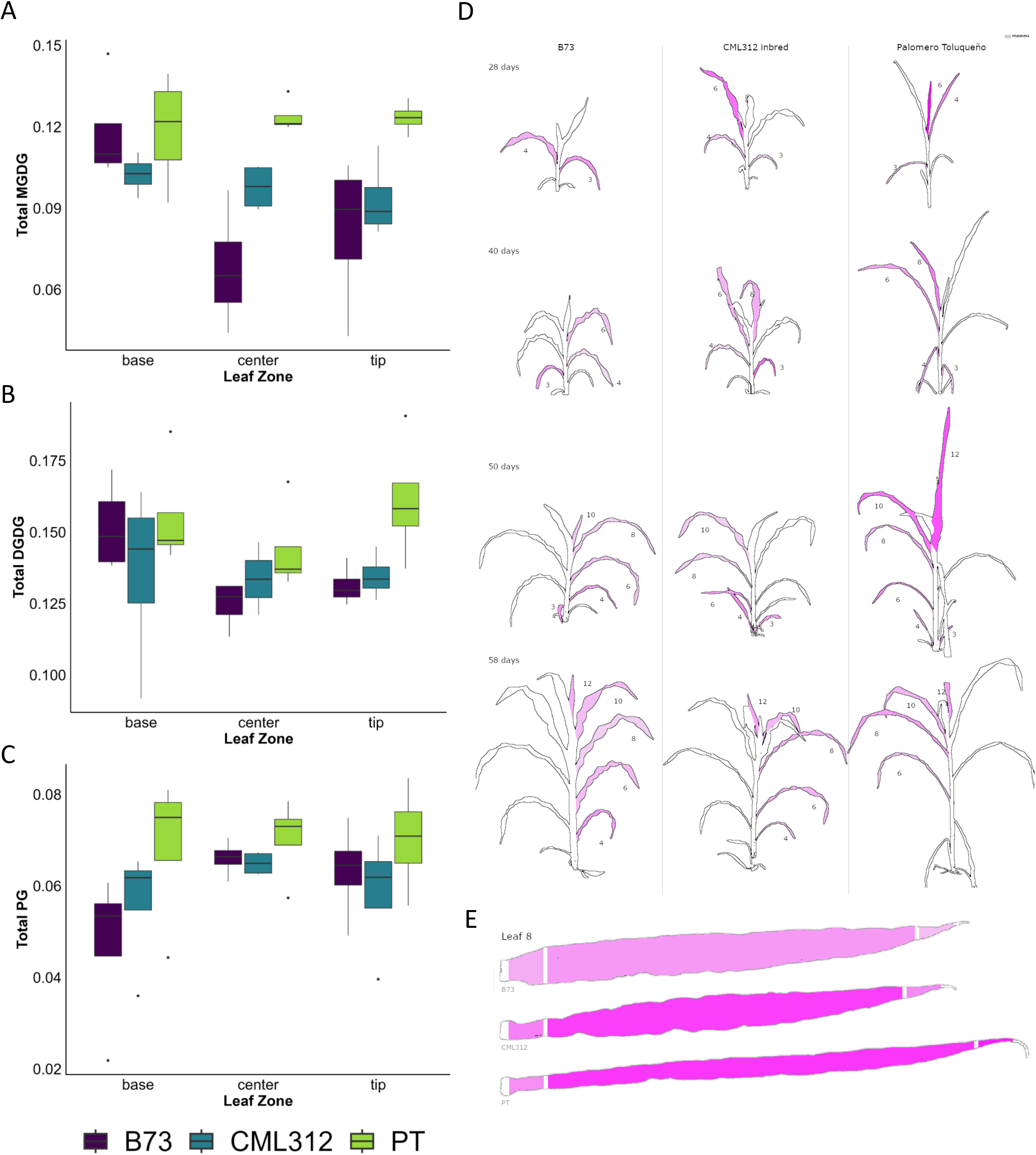
Lipid composition varies by segment of leaf sampled. Monogalactosyldiacylglycerol (MGDG) **(A)**, digalactosyldiacylglycerol (DGDG) **(B)**, and phosphatidylglycerol (PG) **(C)** are examples of lipids that had at least one species with significant variation due to zone as a factor in stepwise modeling. MGDG 36:3 content is shown by genotype, leaf number, and stage of plant **(D)** as well as leaf zone **(E)** in the eFP browser.

### Data Resources for the Community

With this data, we have created a Shiny App to distribute the data from this study and an eFP browser for visualizing the data. The Shiny App allows users to choose a single lipid to examine, view lipid class sums, and filter by specific leaf numbers, genotypes, or leaf regions. The app also displays the images shown in Figure 1, calculates correlations, performs PCA, and allows users to download data. The eFP browser was inspired by the Arabidopsis lipid map eFP browser generated in Kehelpannala *et al*. (2021). The major difference between the maize and Arabidopsis browsers is that the Arabidopsis browser uses multiple tissues from the plant, whereas the maize browser uses just leaf tissue, but multiple genotypes. Raw files can be accessed via Metabolights at accession MTBLS15840.

## CONCLUSION

Together our results establish a reference for maize lipid diversity across genotypes, leaves and developmental stages. Given the known circadian and environmental regulations of plant lipid metabolism, these profiles represent one moment in a dynamic metabolic cycle. Further sampling across the day-night cycle will be essential to resolve how time, growth stage, and environmental transitions further shape lipid remodeling. It is our hope that the data presented in this manuscript, the Shiny App, and the eFP browser may be incorporated into community sites and made openly available to all.

## ACKNOWLEDGMENTS

The authors would like to acknowledge Andi Kur for her work designing Zea Lip covers and logos. This work is supported by the USDA-ARS and the Science and Technologies for Phosphorus Sustainability (STEPS) Center, a National Science Foundation Science and Technology Center (CBET-2019435). NT was supported by the U.S. Department of Energy, Office of Science, Biological and Environmental Research program, Early Career Award Number DE-SC0021889 awarded to RRÁ. NJP was funded by a Discovery Grant from the Natural Sciences and Engineering Research Council of Canada (NSERC), and funding from Genome Canada/Ontario Genomics (OGI-211) supported BAR server upgrades. ACB was supported by NSF-Plant Genome Research Program, Postdoctoral Fellowships in Biology Grant Number 2010703. This work was also supported by the Research Capacity Fund (HATCH), project award no. 7005660, from the U.S. Department of Agriculture’s National Institute of Food and Agriculture, Conacyt Young Investigator (CB-238101), Consejo Nacional de Ciencia y Tecnología National Problems (APN-2983), University of California-MEXUS, and North Carolina State startup funds awarded to RRÁ.

