## Supplemental Figures for "*Zea Lip*: An atlas of glycerolipid profiles across leaf development in maize"

### Slide 1
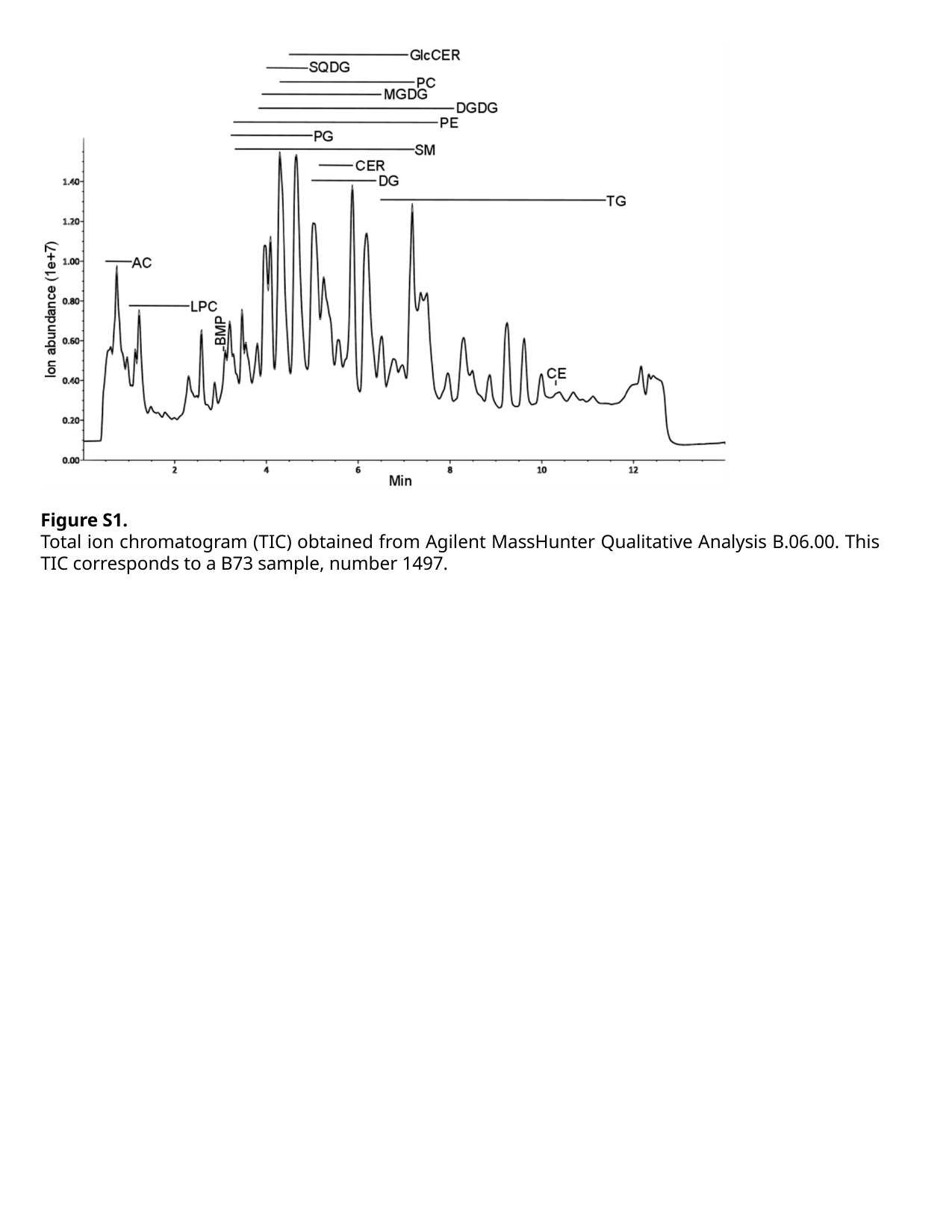

Figure S1.
Total ion chromatogram (TIC) obtained from Agilent MassHunter Qualitative Analysis B.06.00. This TIC corresponds to a B73 sample, number 1497.

### Slide 2
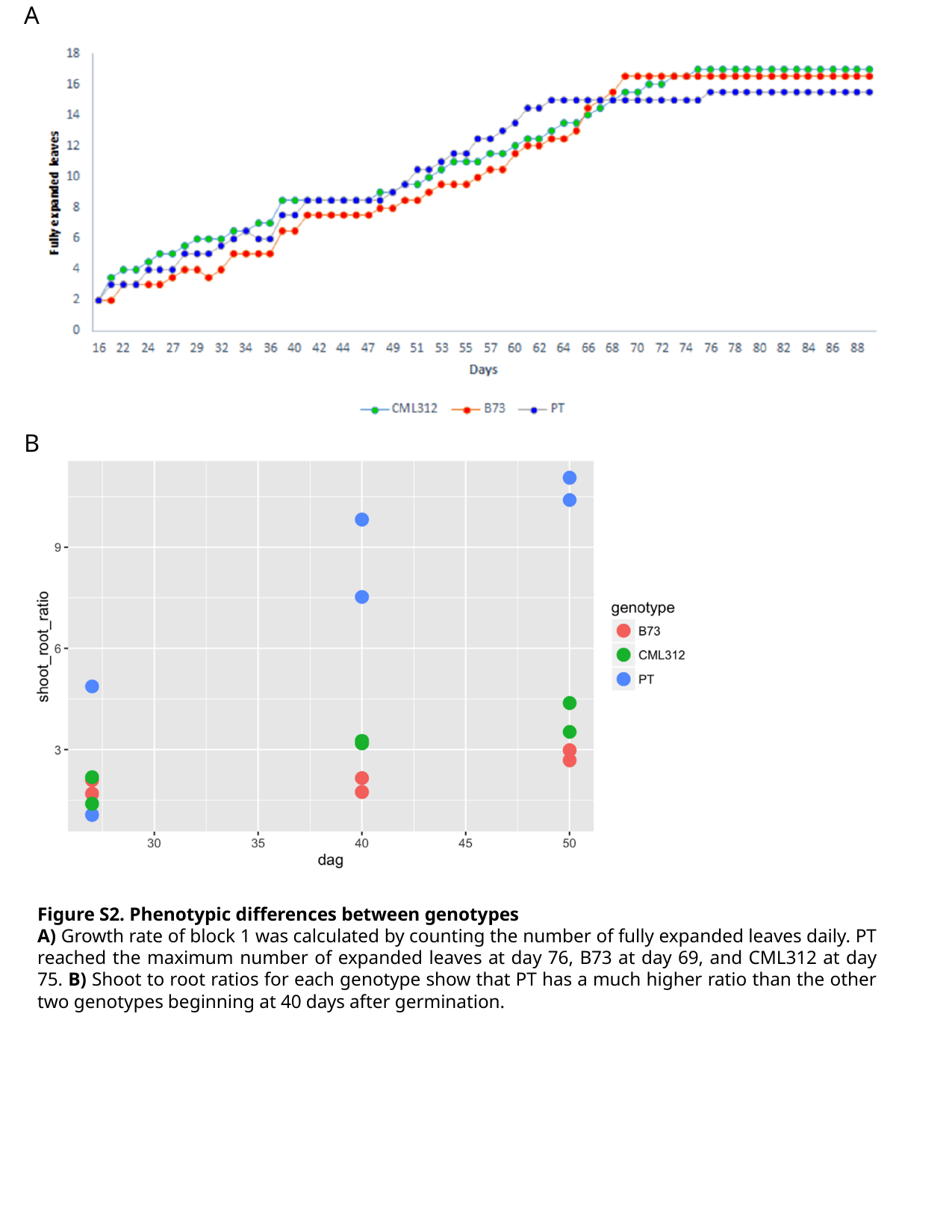

A
B
Figure S2. Phenotypic differences between genotypes
A) Growth rate of block 1 was calculated by counting the number of fully expanded leaves daily. PT reached the maximum number of expanded leaves at day 76, B73 at day 69, and CML312 at day 75. B) Shoot to root ratios for each genotype show that PT has a much higher ratio than the other two genotypes beginning at 40 days after germination.

### Slide 3
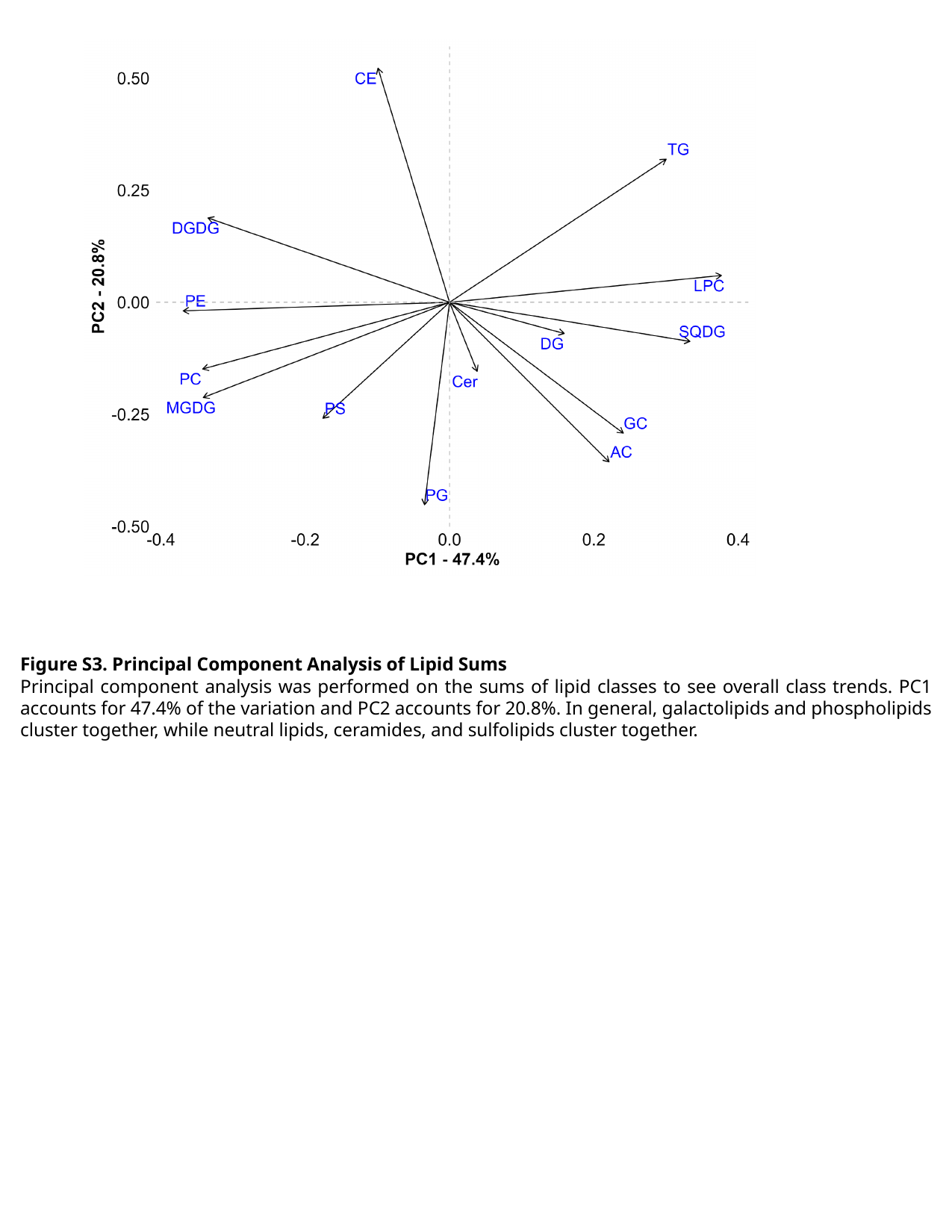

Figure S3. Principal Component Analysis of Lipid Sums
Principal component analysis was performed on the sums of lipid classes to see overall class trends. PC1 accounts for 47.4% of the variation and PC2 accounts for 20.8%. In general, galactolipids and phospholipids cluster together, while neutral lipids, ceramides, and sulfolipids cluster together.

### Slide 4
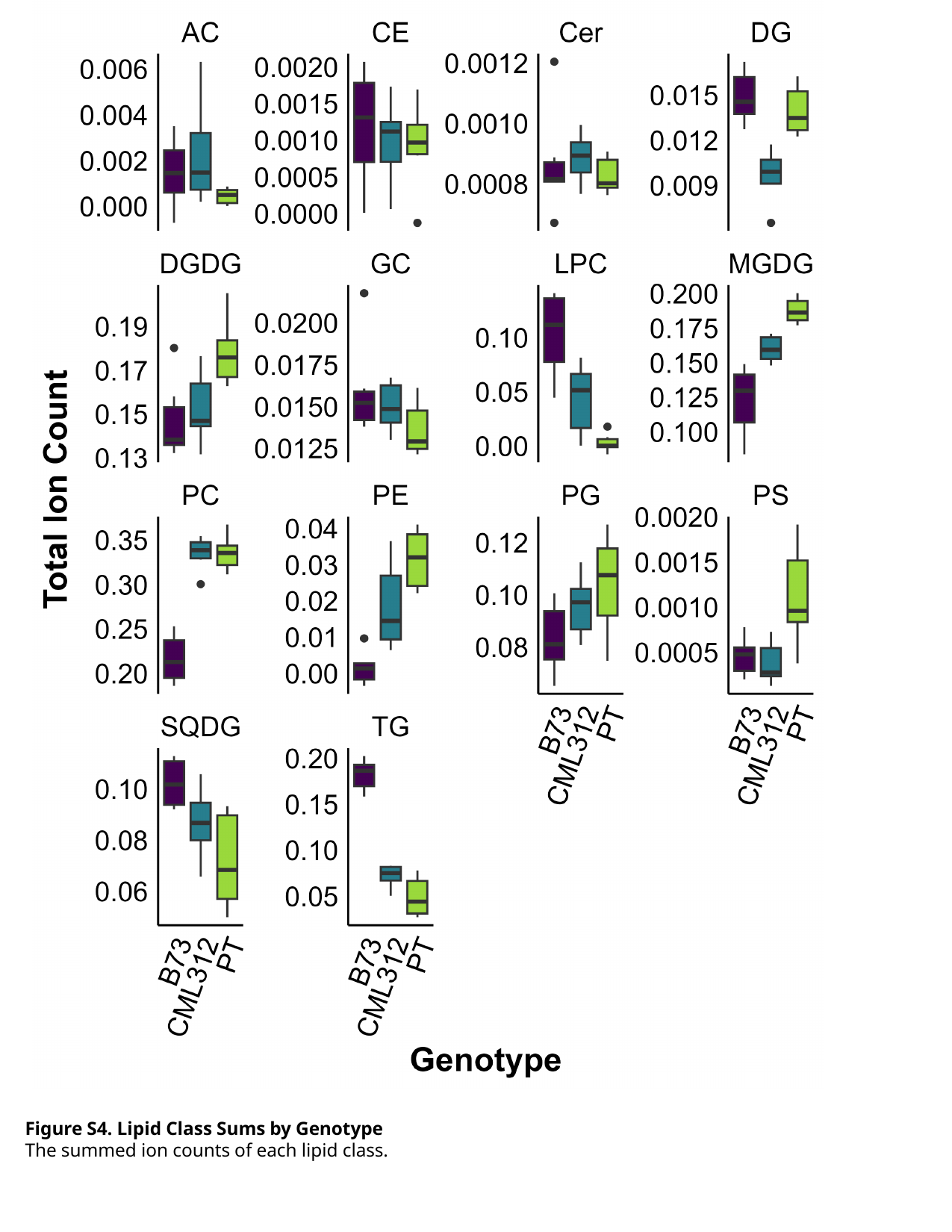

Figure S4. Lipid Class Sums by Genotype
The summed ion counts of each lipid class.

### Slide 5
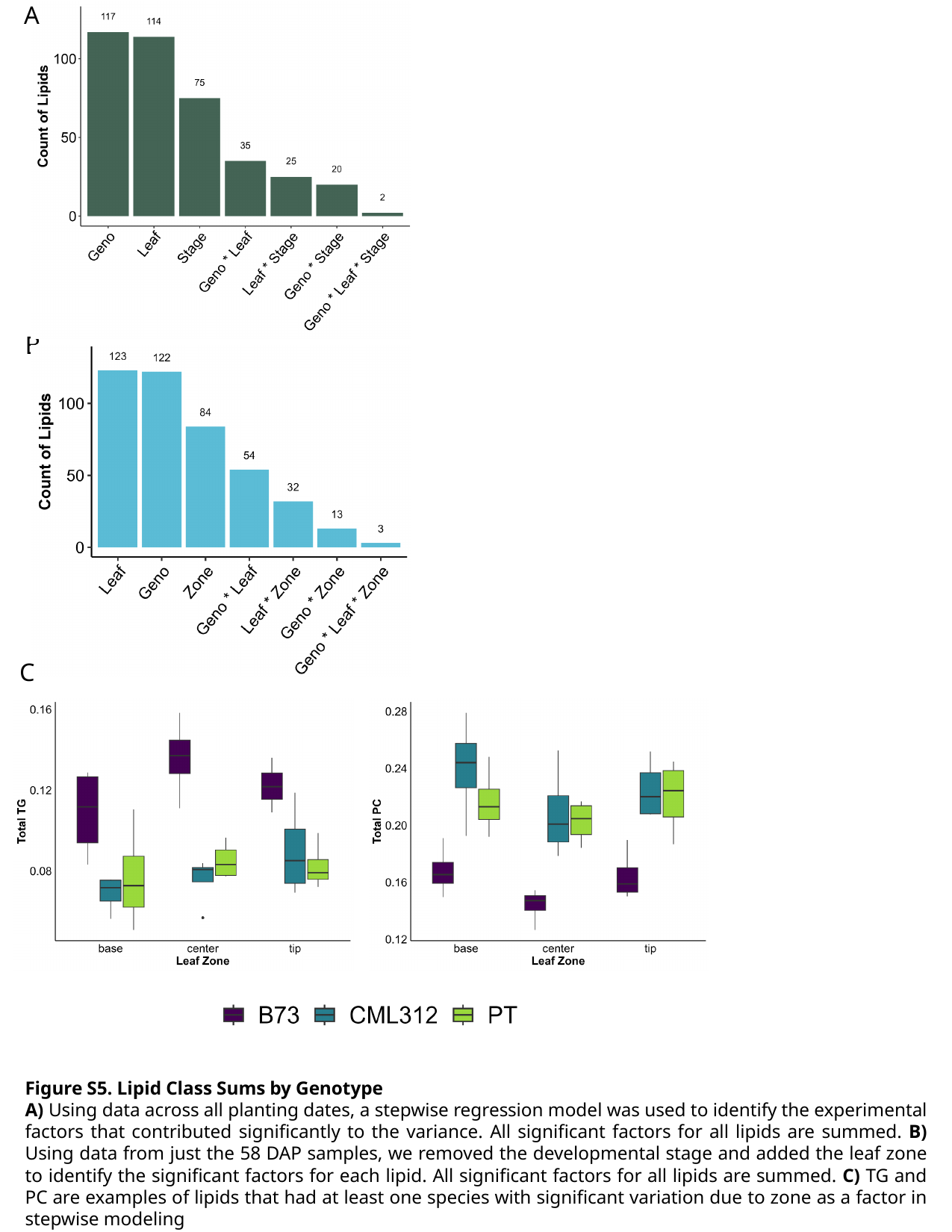

A
C
B
Figure S5. Lipid Class Sums by Genotype
A) Using data across all planting dates, a stepwise regression model was used to identify the experimental factors that contributed significantly to the variance. All significant factors for all lipids are summed. B) Using data from just the 58 DAP samples, we removed the developmental stage and added the leaf zone to identify the significant factors for each lipid. All significant factors for all lipids are summed. C) TG and PC are examples of lipids that had at least one species with significant variation due to zone as a factor in stepwise modeling
